# Microglial reactivity in the hippocampal CA2 is associated with advanced neuronal α-synucleinopathy

**DOI:** 10.64898/2026.02.06.704430

**Authors:** Esteban Luna, Katheryn A. Q. Cousins, Sheina Emrani, Sharon X. Xie, Winifred Trotman, Daniel Weintraub, Alice S. Chen-Plotkin, Edward B. Lee, David J. Irwin

## Abstract

Lewy body diseases are thought to evolve by the spread of intraneuronal α-synuclein pathology. However, staging models of α-synuclein pathology in the human brain seldom evaluate regions with direct synaptic connectivity to model microglial processes in disease. Here, we address this gap by testing the hypothesis that, within the well-defined synaptic connectivity of the intrahippocampal circuit, neuronal α-synuclein pathology is associated with microglial reactivity.

We selected a cohort of autopsy-confirmed Lewy body disease patients with hippocampal neuronal α-synuclein pathology and minimal age-related co-pathologies (*n*=62) and as a control for other intraneuronal hippocampal pathology, we used a cohort of cognitively healthy patients with focal hippocampal tau accumulation with minimal amyloid plaques, termed primary age-related tauopathy (*n*=12). We immunostained consecutive hippocampal sections for α-synuclein pathology and established markers of microglial reactivity, Iba1, HLA-DR, and CD68. With validated digital histology methods, we measured percent area occupied in 6 hippocampal subfields to compare the percent area occupied within subfields between patient cohorts, correlate the percent area occupied within and between subfields, and use linear-mixed effects models to compare the percent area occupied across subfields while co-varying for demographics. We also constructed two groups of either low-level hippocampal α-synuclein pathology restricted to the cornu ammonis 2-3 subfields (Focal Subtype) or widespread α-synuclein pathology within additional subfields (Widespread Subtype) to model hypothesized α-synuclein pathological spread within the intrahippocampal circuit.

Lewy body disease patients exhibited increased HLA-DR and CD68 in most hippocampal subfields compared with primary age-related tauopathy. In Lewy body disease patients, staining for all microglial markers was highest in the cornu ammonis field 2. Neuronal α-synuclein pathology in the cornu ammonis field 2 consistently correlated with only HLA-DR and CD68 but not Iba1. Patients classified as the Widespread Subtype had worse cognitive impairment and increased HLA-DR and CD68 within the cornu ammonis field 2. Furthermore, HLA-DR and CD68 in the cornu ammonis field 2 correlated with distal neuronal α-synuclein pathology only in subfields with retrograde connectivity.

We found increased hippocampal microglial abnormalities in Lewy body disease patients compared with cognitively normal controls with primary age-related tauopathy, suggesting a relatively specific microglial/inflammatory response to neuronal α-synuclein pathology. Our data is consistent with predominantly retrograde transmission of neuronal α-synuclein pathology across hippocampal subfields, where the focal microglial response in the cornu ammonis field 2 may influence clinical outcomes and α-synuclein pathological spread. These data suggest that microglial processes can help refine Lewy body disease histopathological staging.

## Introduction

Intraneuronal protein aggregates predominantly made up of α-synuclein, termed neuronal α-synuclein pathology (n-asyn), pathologically define Lewy body diseases (LBDs) that clinically encompasses Parkinson’s disease (PD) without and with dementia (PDD) and dementia with Lewy bodies.^1–3^ Braak and colleagues initially proposed a topographical pattern of n-asyn split into progressive stages in PD from the olfactory bulb and dorsal medulla with hypothesized spreading rostrally from the brainstem to limbic and neocortical regions.^4,5^ Subsequent postmortem staging systems define additional patterns of progressive n-asyn transsynaptic spread in the human brain and peripheral nervous system, but thus far, these schemata do not account for non-neuronal aspects of disease pathophysiology or provide detailed analysis within the anatomic framework of local circuits to model these intercellular interactions.^6–12^ Indeed, genome-wide association studies in LBD have found risk variants in microglia-associated genes, and postmortem studies have found significant morphologic, transcriptional, and molecular changes in microglia within specific disease relevant brain regions, such as the Substantia nigra pars compacta, suggesting a microglial component to disease pathogenesis.^13–17^ Limited studies suggest relatively modest increased microglial reactivity to n-asyn compared to the microglial response in Alzheimer’s disease (AD).^18–23^ Still the relationship between LBD clinical progression, increasing n-asyn burden within key vulnerable brain regions, and microglial reactivity in LBD is unclear. The hippocampus provides an ideal anatomic model to test the association of microglial reactivity to progressive n-asyn burden in LBD because the subfields form part of the well-defined intrahippocampal circuit that is critical to memory encoding and affected in LBD.^24,25^ The hippocampus develops n-asyn in dementia with Lewy bodies and in stage V of classical Braak staging of PD/PDD.^4,26^ The cornu ammonis 2 (CA2) hippocampal subfield is particularly vulnerable to forming n-asyn compared to other hippocampal subfields in LBD.^26–28^ Moreover, one study found that the CA2 develops increasing microglial phagocytic activity with increasing Parkinson’s Braak stage by measuring staining to CD68, but detailed anatomic studies to model the relationship of microglial reactivity with subregional n-asyn burden in the hippocampus are limited.^21^ This is a crucial knowledge gap that could provide critical clues in elucidating the role of microglia in LBD pathogenesis.

Here, we tested the hypothesis that microglial reactivity is associated with n-asyn burden within the hippocampal subfields in LBD. To do this, we selected a unique cohort of LBD patients with relatively pure n-asyn neuropathology and limited potentially confounding age-related co-pathologies.^29,30^ To establish if microglial reactivity was specific to n-asyn and not to other age-related intraneuronal pathology, we compared this select LBD cohort to a well-characterized control cohort of healthy adults diagnosed post-mortem with primary age-related tauopathy (PART), defined as limited age-related tau pathology within the CA2.^31,32^ We used rigorously validated digital histopathology methods to measure the percent area occupied (%AO) of n-asyn and classically associated pro-inflammatory microglial markers of total microglial area, phagocytic activity, and antigen presentation with Iba1, CD68, and HLA-DR, respectively.^28,33–36^ We found that LBD patients exhibit greater CD68 and HLA-DR signal in almost all hippocampal subregions compared to PART controls without increased microglial total area measured by Iba1 arguing for increased microglial reactivity specific to n-asyn without a greater infiltration of microglia into the hippocampus. Moreover, we found that LBD patients with widespread hippocampal n-asyn throughout the hippocampal subfields compared to those with limited hippocampal n-asyn in the CA2-3 subfields alone demonstrate increased focal microglial reactivity within the CA2 and worse cognition. This focal CA2 microglial reactivity correlated with n-asyn burden in distal subfields only with retrograde connectivity to CA2-3 region within the intrahippocampal circuit. Contrary to our hypothesis, the data presented here implicate only localized CA2 microglial reactivity to n-asyn in both advancing LBD clinical progression and neuropathological progression within the hippocampus. Furthermore, these data provide evidence that deeper phenotyping beyond n-asyn localization alone can further refine neuropathologic staging of LBD subtypes.

## Materials and methods

### Patient selection and study workflow

We selected patient samples from the Penn Integrated Neurodegenerative Disease Database (INDD) on 09/26/22 with the following criteria: To maximize the number of samples with n-asyn within the hippocampus, we first had primary inclusion criteria of a clinical diagnosis of PD, PDD, or dementia with Lewy bodies) and primary neuropathologic diagnosis of LBD (i.e. brainstem, limbic or neocortical stage, *n*= 351). To reduce the confounds of mixed pathology, we excluded cases that had a clinically relevant medium- or high-level AD neuropathologic change (*n*=154 met inclusion criteria).^37^ We selected available tissue from the hippocampus at the level of the lateral geniculate nucleus for optimal sampling of the 6 hippocampal subfields (CA 1-4, dentate gyrus [DG], subiculum [Sub]). Thus, we excluded cases with missing/damaged tissues and/or suboptimal sampling of the hippocampal subfields at this coronal level, resulting in a final cohort of 62 patients (Fig. 1A). We included a control group of healthy adult brain donors without clinical neurologic disease, with a primary neuropathologic diagnosis of PART, minimal other age-related pathology, and available hippocampal tissue as above (*n*=12).

**Figure 1.**
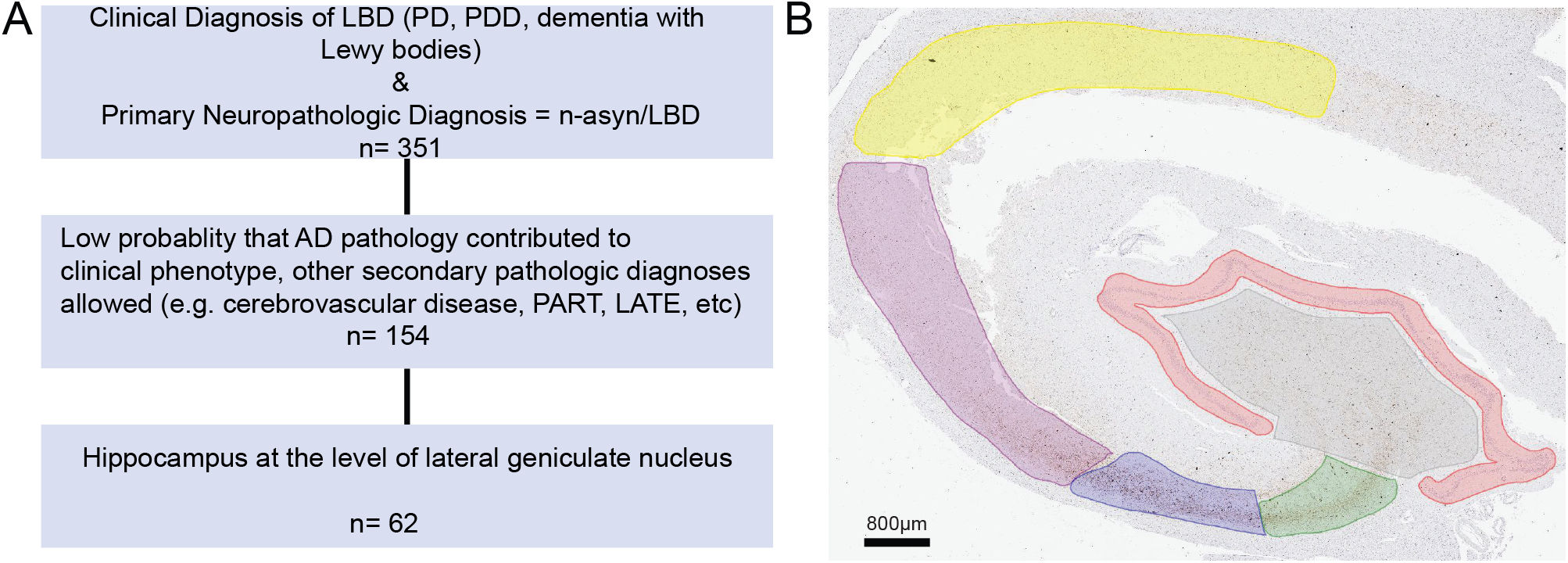
Study design and hippocampal annotation. (**A**) The flow diagram describes the inclusion and exclusion criteria for selecting patient samples for the LBD cohort. (**B**) Hippocampal subfield annotations of a LBD patient sample stained for n-asyn. Red = DG, Grey = CA4, Green = CA3, Blue = CA2, Maroon = CA1, Yellow = Sub. Scale bar indicated in image.

### Neuropathologic assessment

Diagnostic neuropathologic assessment was performed as previously described using current neuropathological criteria for AD and LBD.^37,38^ Briefly, fresh brains obtained at the time of autopsy were fixed in either neutral buffered formalin or 70% ethanol overnight. The brain tissue was processed and embedded in paraffin as previously described.^38^ 6μm thick sections were stained by immunohistochemistry (IHC) with validated antibodies to tau, amyloid-β, n-asyn, and TDP-43. All protocols were performed with informed consent and compliant with the University of Pennsylvania Institutional Review Board (IRB).

### Digital neuropathologic assessment

Near adjacent tissue sections were stained by IHC in the Penn Digital Neuropathology Lab for phosphorylated-α-synuclein S129 as a marker of n-asyn (Abcam, MJF-R13, ab209421, 1:20k), Iba1 (FUJIFILM Wako, 019-19741, 1:4k, antigen retrieval with citric acid for 15 minutes), CD68 (Dako, M081401, 1:100, antigen retrieval with citric acid for 15 minutes), and HLA-DR (Genetex, MSVA-478R, GTX04457, 1:100, antigen retrieval with citric acid for 15 minutes) as previously described.^39,40^ Stained IHC sections were digitally scanned (Aperio AT2, Leica Biosystem, Wetzlar, Germany) at 20x magnification. Hippocampal subfield annotation and digital histology analysis was performed using QuPath Software (version 0.5.1) as previously described using previously validated cytoarchitectonic definitions of subfield delineations to generate a %AO of immunostain signal for each antibody in each region of interest (ROI; CA1-4, DG, Sub) from each whole-slide image of a unilateral hippocampal section for each patient that met all inclusion criteria.^41,42^ An illustrative example of hippocampal subfield annotation to measure CA1-4, DG, and Sub is shown (Fig. 1B).

### Clinical assessment

Clinical assessment of patients was done as part of ongoing observational research studies at Penn (P01-AG084497, P30-AG072979). These clinical assessments have been previously described.^43^ Due to changes in batteries over the time-period of autopsies, we focused on the Montreal Cognitive Assessment (MOCA) and Dementia Rating Scale-2 (DRS), which had the greatest amount of harmonized data and are relevant scores to memory. For patients with longitudinal data, the assessment closest to death was used to relate to postmortem data.

### n-asyn hippocampal staging

To model progressive n-asyn spread through the intrahippocampal circuit, we dichotomized patients into 2 groups or stages based on visual inspection. The group, termed Focal Subtype, was identified by n-asyn being relatively localized to the CA2-3 region with no or minimal n-asyn in other subfields. The group, termed Widespread Subtype, was identified as sections with substantial n-asyn distributed in hippocampal subfields beyond the CA2-3 region. Dichotomization was done by a single rater using solely the n-asyn-stained slides for grouping. There were no samples with n-asyn present predominantly outside of the CA2-3 region in this LBD cohort.

### Statistical analysis

Group comparisons of demographics and clinical data were performed using independent sample two-tailed t-tests for continuous measures that follow normality assumptions or Wilcoxon signed-rank test for non-normally distributed continuous measures where indicated. Correlations of %AO were done using non-parametric spearman rank correlation. To compare overall patterns of regional burden of immunostain %AO, we used linear mixed-effects models, which account for correlations among multiple sample data points within an individual. %AO values were assessed for normality and transformed using natural log. Natural log %AO was used as the dependent variable with fixed effect independent variables of age, sex, post-mortem interval (PMI) in hours, race (white, non-white, unknown), Consortium to Establish a Registry for Alzheimer’s Disease (CERAD) plaque score (C0, C1, C2, C3), Thal Amyloid Stage (A0, A1, A2, A3), Alzheimer’s tau Braak Stage (B0, B1, B2, B3), clinical syndrome (PD, PDD, dementia with Lewy bodies) and hippocampal subfield ROI as a fixed factor with the CA2 as the reference strata to identify regional patterns of disease.^44,45^ A random intercept term was included in mixed-effects models. All statistical analyses were performed using Rstudio (Version 2024.04.2+764). Data is presented using mean ± standard deviation (SD) unless otherwise indicated with a threshold of *p*=0.05 for significance based on the hypothesis-driven nature of our regional comparisons.

## Results

### LBD has increased hippocampal microglial reactivity relative to PART

We first set out to test if microglial reactivity in LBD hippocampal subfields is specifically associated with n-asyn in this relatively “pure” LBD cohort to limit the effect of amyloid and tau co-pathology on microglial measures.^29,30^ We did this by comparing the microglial reactivity in the hippocampal subfields with a control PART cohort without antemortem cognitive deficits. A summary of demographics, neuropathologic findings (including presence of low levels of other age-related pathologies), and clinical data is summarized in Table 1. Briefly, the cohorts were similar in demographics except age, where the PART cohort was younger, with a mean age at death of 64 (Range: 51-83) compared to the LBD cohort who had a mean age at death of 78 (Range: 58-98) (*p*<0.001). We found that LBD and PART cohorts had similar staining %AO for the general microglial marker Iba1 in all hippocampal subfields (Fig. 2A, B).^33^ We evaluated for reactivity to the microglial markers of phagocytic activity and antigen presentation by evaluating for CD68 and HLA-DR %AO, respectively, in the hippocampal subfields. There was increased HLA-DR %AO in LBD compared to PART in all subfields except the CA4 (Fig. 2A, C). CD68%AO was also increased in LBD compared to the PART in all subfields except the DG (Fig. 2A, D). These data illustrate that despite both LBD and PART being characterized by intraneuronal protein accumulation in the hippocampus, especially in the CA2 subfield, LBD patients selectively exhibit increased measures of microglial reactivity to neuropathology in most hippocampal subfields without overt overall microglial infiltration marked by Iba1.^33,36^

**Table 1.**
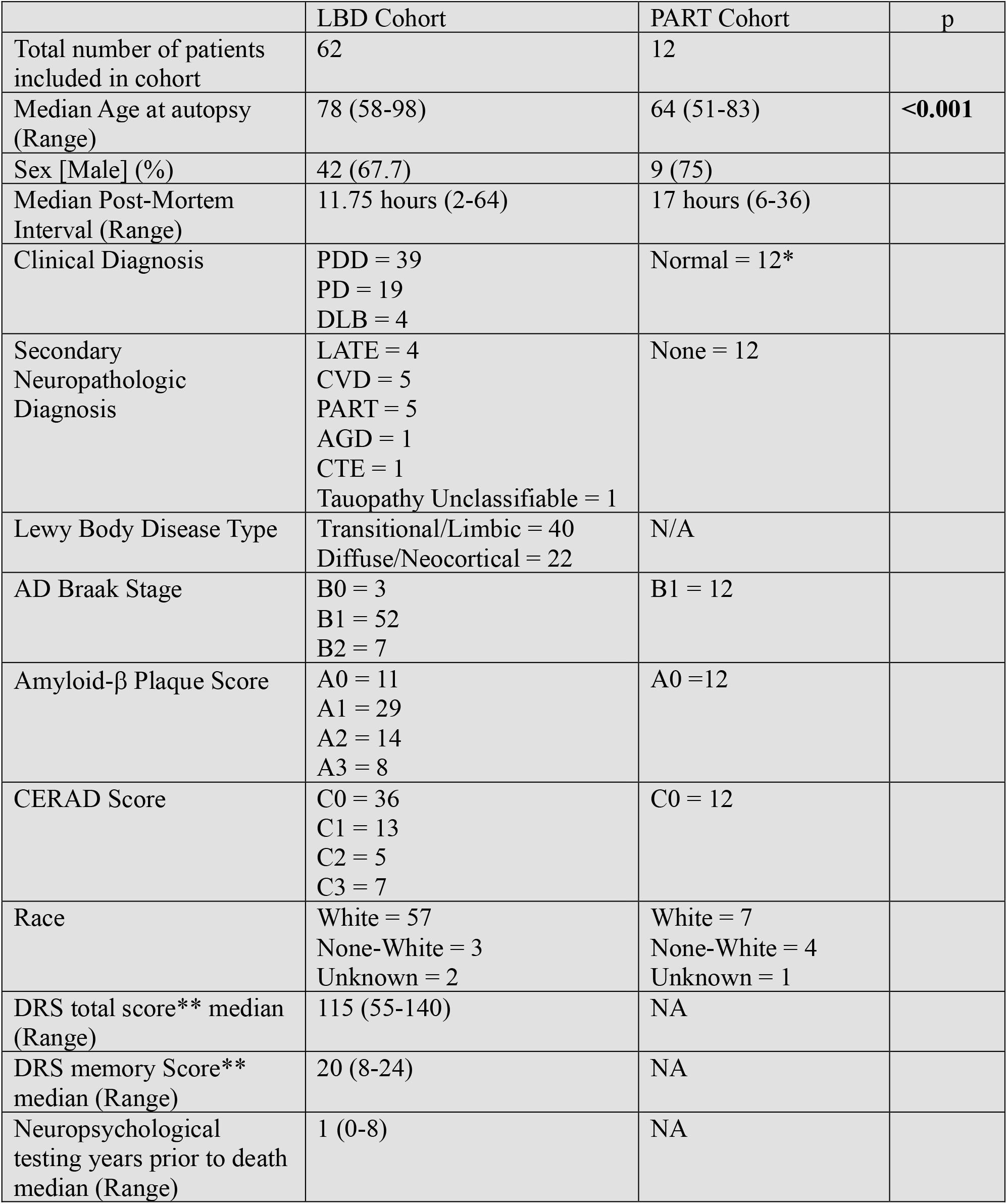
Patient demographic and pathological characteristics. Abbreviations. PDD = Parkinson’s disease with Dementia, PD = Parkinson’s disease, DLB = Dementia with Lewy Bodies, AD = Alzheimer’s disease, LATE = Limbic-predominant age-related TDP-43 associated encephalopathy, PART = Primary age-related tauopathy, CTE = Chronic traumatic encephalopathy, DRS = Dementia Rating Scale. NA= not available * one patient had a primary psychiatric diagnosis but no neurodegenerative dementia. ** DRS scores were only available for 28 patients

|  | LBD Cohort | PART Cohort | p |
| --- | --- | --- | --- |
| Total number of patients included in cohort | 62 | 12 |  |
| Median Age at autopsy (Range) | 78 (58-98) | 64 (51-83) | **<0.001** |
| Sex [Male] (%) | 42 (67.7) | 9 (75) |  |
| Median Post-Mortem Interval (Range) | 11.75 hours (2-64) | 17 hours (6-36) |  |
| Clinical Diagnosis | PDD = 39<br>PD = 19<br>DLB = 4 | Normal = 12* |  |
| Secondary Neuropathologic Diagnosis | LATE = 4<br>CVD = 5<br>PART = 5<br>AGD = 1<br>CTE = 1<br>Tauopathy Unclassifiable = 1 | None = 12 |  |
| Lewy Body Disease Type | Transitional/Limbic = 40<br>Diffuse/Neocortical = 22 | N/A |  |
| AD Braak Stage | B0 = 3<br>B1 = 52<br>B2 = 7 | B1 = 12 |  |
| Amyloid-β Plaque Score | A0 = 11<br>A1 = 29<br>A2 = 14<br>A3 = 8 | A0 = 12 | |
| CERAD Score | C0 = 36<br>C1 = 13<br>C2 = 5<br>C3 = 7 | C0 = 12 |  |
| Race | White = 57<br>None-White = 3<br>Unknown = 2 | White = 7<br>None-White = 4<br>Unknown = 1 |  |
| DRS total score** median (Range) | 115 (55-140) | NA |  |
| DRS memory Score** median (Range) | 20 (8-24) | NA |  |
| Neuropsychological testing years prior to death median (Range) | 1 (0-8) | NA |  |

**Figure 2.**
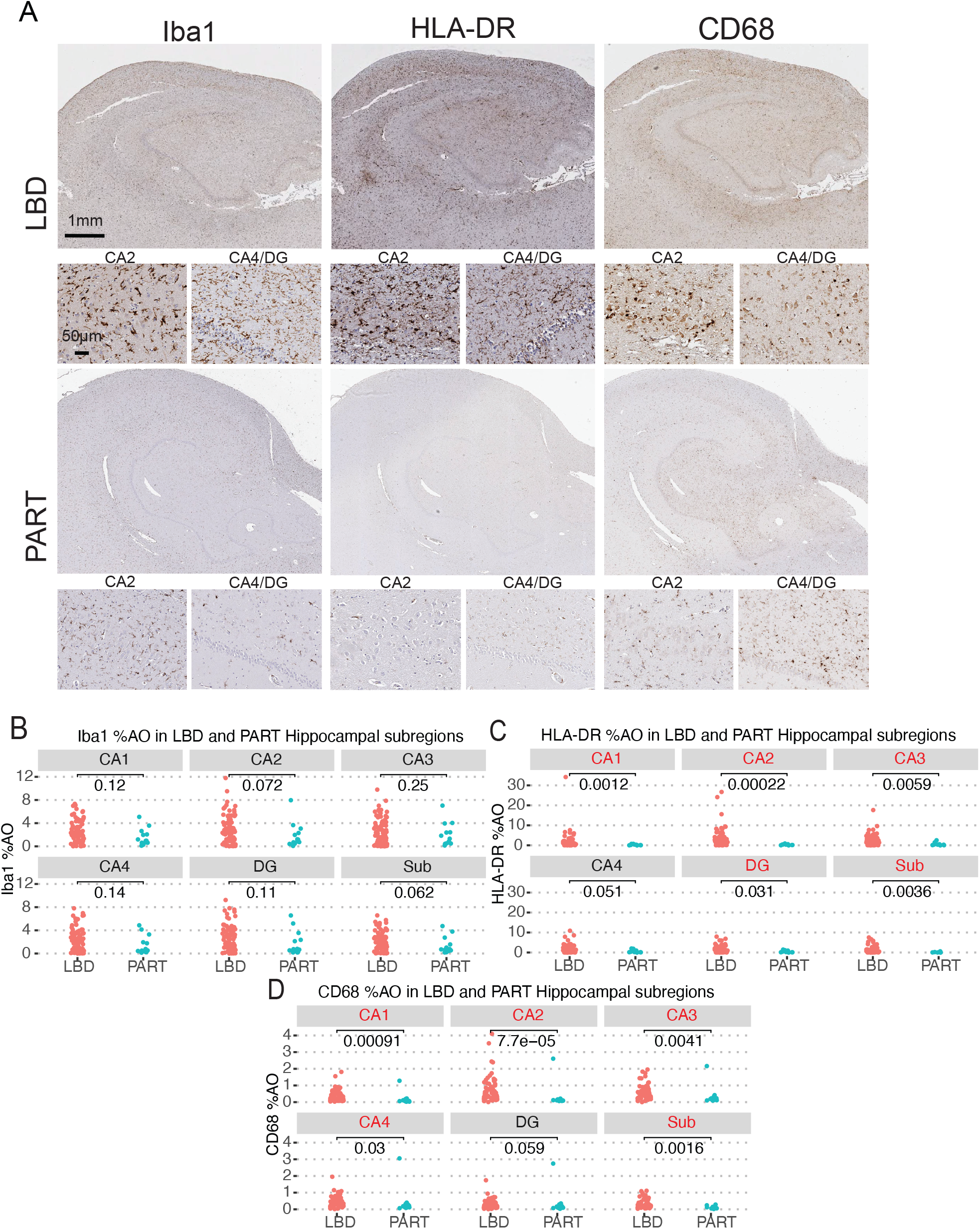
Hippocampal microglial reactivity is increased in LBD relative to PART controls. (**A**) Near adjacent hippocampal sections stained for Iba1, HLA-DR, and CD68 from a LBD patient and a PART patient. Insets from CA2 and CA4/DG from each image. Scale bars are indicated in image. (**B**) Comparison of %AO of Iba1, (**C**) %AO HLA-DR and (**D**) %AO CD68 between PART and LBD cohorts in each hippocampal subregion. All Statistical analyses done using Wilcoxon-signed rank test for non-parametric data. Each data point represents 1 patient sample for that subregion. *n*=11 for PART cohort in Sub due to damaged tissue precluding measurement. *p*-values for each comparison is provided on graphs. All statistically significant results demarcated in red font of graph title.

### The CA2 in LBD shows the strongest correlation between n-asyn and microglial reactivity

To determine the regional distribution and potential association of microglial reactivity to n-asyn, we tested the regional severity of n-asyn and the microglial markers among the hippocampal subfields in the LBD cohort using linear mixed effects modeling while adjusting for demographic variables (Table 2). Similar to previous findings, we found that the CA2 had the greatest n-asyn levels (Fig. 3A; Table 2).^27^ For the general microglial marker, Iba1, we found higher %AO in the CA2 compared with the Sub, CA4, and DG, but not the CA1 or 3 (Fig. 3B; Table 2). HLA-DR showed the highest %AO in the CA2 relative to all other subregions (Fig. 3C, Table 2). CD68 also showed the highest %AO in the CA2 compared with all other hippocampal subregions except the CA3 which had similarly high levels (Fig. 3D, Table 2). Interestingly, examination of demographic predictors in our models found that male sex associated with increased HLA-DR %AO and CD68 %AO and that white patients demonstrated higher CD68 %AO compared to none-white LBD patients (Table 2).

**Table 2.** Linear mixed effect model for distribution of Iba1, HLA-DR and CD68 in LBD cohort hippocampus. *CA2 is the reference category. ** DLB is the reference category. ***None-white patients is the reference category. All coefficients for the variables reported in comparison to the reference category.

| Stain | Iba1 |  |  | HLA-DR |  |  | CD68 |  |  |
| --- | --- | --- | --- | --- | --- | --- | --- | --- | --- |
| Predictors | $\beta$ | SE | p | $\beta$ | SE | p | $\beta$ | SE | p |
| Region* |  |  |  |  |  |  |  |  |  |
| Subiculum | -0.22 | 0.08 | **<0.01** | -0.99 | 0.08 | **<0.001** | -0.62 | 0.76 | **<0.001** |
| CA1 | -0.14 | 0.08 | 0.08 | -0.73 | 0.08 | **<0.001** | -0.53 | 0.76 | **<0.001** |
| CA3 | -0.09 | 0.08 | 0.27 | -0.24 | 0.08 | **<0.01** | -0.10 | 0.76 | 0.19 |
| CA4 | -0.32 | 0.08 | **<0.001** | -0.62 | 0.08 | **<0.001** | -0.29 | 0.76 | **<0.001** |
| DG | -0.18 | 0.08 | **<0.05** | -0.72 | 0.08 | **<0.001** | -0.66 | 0.76 | **<0.001** |
| Clinical Subgroup** |  |  |  |  |  |  |  |  |  |
| PD | 0.10 | 1.02 | 0.92 | -0.72 | 0.92 | 0.44 | -0.36 | 0.57 | 0.53 |
| PDD | -0.24 | 0.99 | 0.80 | -0.75 | 0.89 | 0.40 | -0.48 | 0.55 | 0.39 |
| Sex [Male] | -0.01 | 0.41 | 0.99 | 0.82 | 0.37 | **<0.05** | 0.75 | 0.23 | **<0.01** |
| Race*** |  |  |  |  |  |  |  |  |  |
| Unknown | 2.14 | 1.65 | 0.20 | 0.09 | 1.49 | 0.95 | 1.81 | 0.91 | 0.05 |
| White | 0.79 | 0.88 | 0.37 | 0.05 | 0.79 | 0.95 | 1.21 | 0.48 | **<0.01** |
| Tau Braak Stage | 0.24 | 0.51 | 0.63 | 0.48 | 0.46 | 0.30 | 0.12 | 0.28 | 0.67 |
| Aβ Thal Stage | 0.39 | 0.27 | 0.16 | -0.32 | 0.24 | 0.20 | -0.14 | 0.15 | 0.34 |
| CERAD Plaque Score | -0.12 | 0.24 | 0.60 | -0.22 | 0.22 | 0.30 | -0.09 | 0.13 | 0.51 |
| Post mortem Interval | 0.03 | 0.02 | 0.10 | 0.03 | 0.17 | 0.13 | 0.01 | 0.01 | 0.18 |
| LBD Stage | -0.03 | 0.42 | 0.94 | -0.61 | 0.38 | 0.12 | -0.41 | 0.23 | 0.09 |
| (Intercept) | -1.11 | 1.55 | 0.48 | 0.62 | 1.41 | 0.66 | -1.84 | 0.86 | **<0.05** |

**Figure 3.**
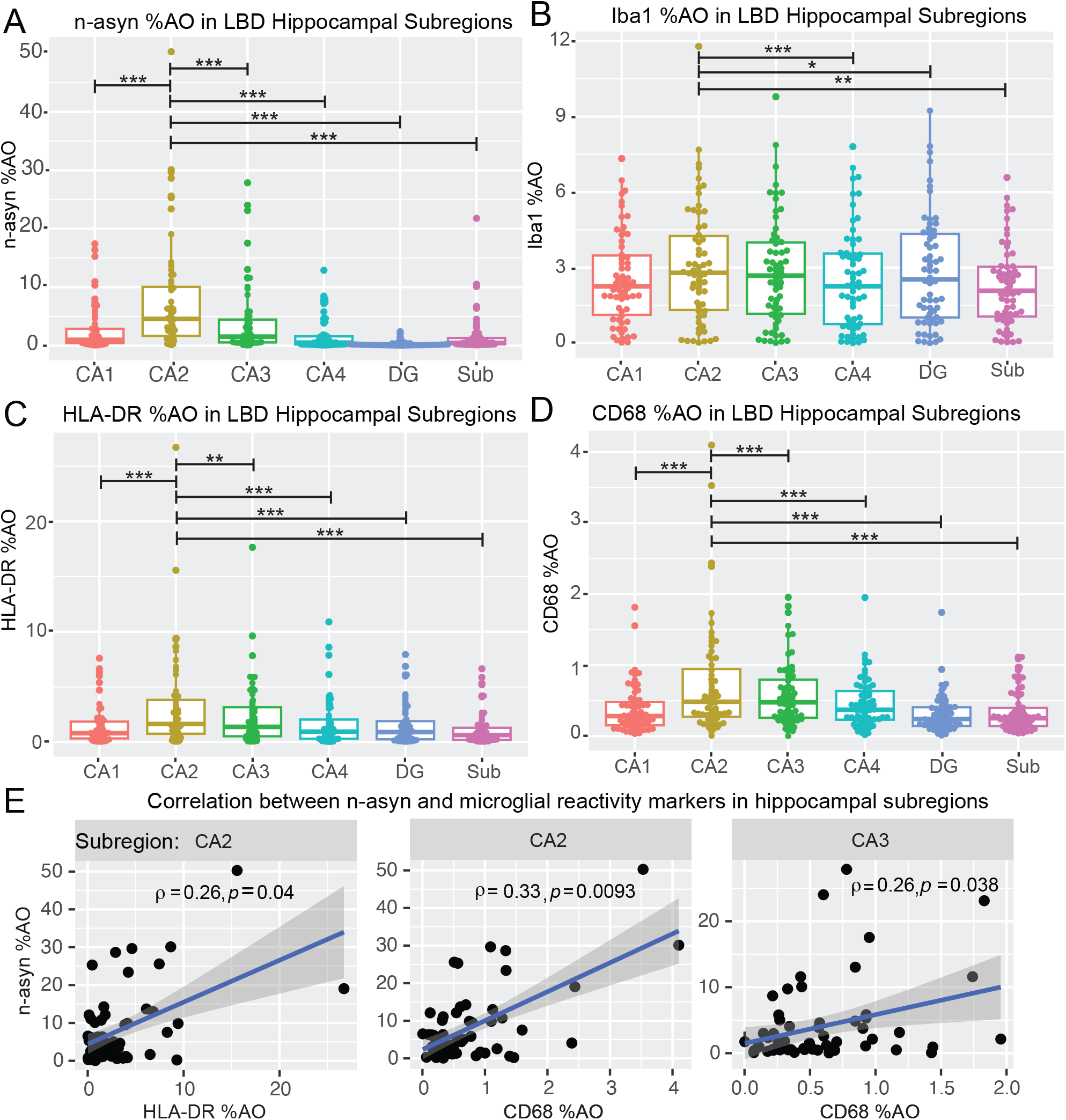
Distribution and correlation of n-asyn and microglial markers in LBD cohort. (**A**) Box plot of n-asyn %AO, (**B**) Iba1 %AO, (**C**) HLA-DR %AO, (**D**) CD68 %AO in LBD cohort. Reported *p*-values are derived from natural-log normalized %AO values in linear mixed effects modelling reported in Table 2. (**E**) Spearman rank correlation in indicated subfields of n-asyn %AO with HLA-DR %AO and CD68 %AO. * *p*<0.05, ** *p*<0.01, *** *p*<0.001. Blue line is best fit line, and dark grey is confidence interval.

To determine if the measured microglial reactivity to n-asyn in the LBD hippocampus mirrors n-asyn burden, we tested correlations of n-asyn with each microglial marker within each hippocampal subfield. n-asyn did not correlate with Iba1 %AO in any hippocampal subregion, arguing against a linear infiltration of microglia in response to n-asyn burden (Supplementary Fig.1). Interestingly, both HLA-DR %AO (*ρ* = 0.26, *p*<0.05) (Fig. 3B) and CD68 %AO (*ρ* = 0.33, *p*<0.01) correlated with n-asyn %AO in the CA2 subfield (Fig. 3E). Within-subfield correlations between n-asyn and HLA-DR didn’t reach significance in any other subfield (Supplementary Fig. 2). CD68%AO additionally correlated with n-asyn %AO in the CA3 (*ρ* = 0.26, *p*<0.05) and the subiculum (*ρ* = 0.3, *p*<0.05) (Fig. 3E; Supplemental Fig. 3). Within subfield correlations between n-asyn and CD68 didn’t reach significance in any other subfield (Supplementary Fig. 3). In summation, the CA2 subfield consistently showed the highest amount of staining for IBA1, CD68, and HLA-DR compared with other subfields, and in contrast to other hippocampal subfields, only the CA2 showed a moderate correlation between n-asyn and CD68 and HLA-DR.

### Greater hippocampal n-asyn spatial distribution is associated with worse clinical outcomes

Braak staging postulates that greater spatial distribution of n-asyn corresponds to a later disease stage and worse clinical outcomes.^4,5^ Thus, we set out to determine if greater spatial distribution of n-asyn within the intrahippocampal circuit was also associated with worse clinical outcomes and increasing n-asyn-induced microglial reactivity. Based on visual inspection, we dichotomized patients into two groups. Those that exhibit n-asyn confined within the CA2-3 regions, classified as Focal Subtype, and patients where additional n-asyn was present outside of the CA2-3 region, classified as Widespread Subtype (Fig. 4A). To ensure that subtyping yielded dichotomized groups of n-asyn burden, we compared n-asyn burden in each subfield between these 2 groups and found that the Widespread Subtype exhibited increased n-asyn in all subregions providing neuropathological confirmation our groupings (Supplemental Fig. 4A-F). Because it is well-established that clinical progression is proportional to n-asyn distribution patterns observed postmortem, we hypothesized that pathology distribution outside of the CA2-3 would correspond to worse clinical outcomes and provide clinical validation of our patient groupings.^4–6^ Within the subset of LBD patients with antemortem DRS scores (*n*=19 Focal Subtype; *n*=9 Widespread Subtype), we found that Widespread Subtype patients had a lower DRS total score (Widespread Subtype: 97.11 ± 20.50; Focal Subtype: 119.16 ± 16.78; *p*=0.006) and DRS memory score (Widespread Subtype: 15.11 ± 4.96; Focal Subtype: 19.63 ± 3.73; *p*=0.012) compared with the Focal Subtype group (Fig. 4B, C). We also found a non-significant trend for lower MoCA scores (*n*=17 Focal Subtype; *n*=8 Widespread Subtype) in Widespread Subtype compared to Focal Subtype patients (Widespread Subtype: 14.75 ± 5.06; Focal Subtype: 18.59 ± 4.01; *p*=0.052; Supplemental Fig. 5). These data imply that greater spatial distribution of n-asyn within the intrahippocampal circuit generally correlates with worse clinical progression.

**Figure 4.**
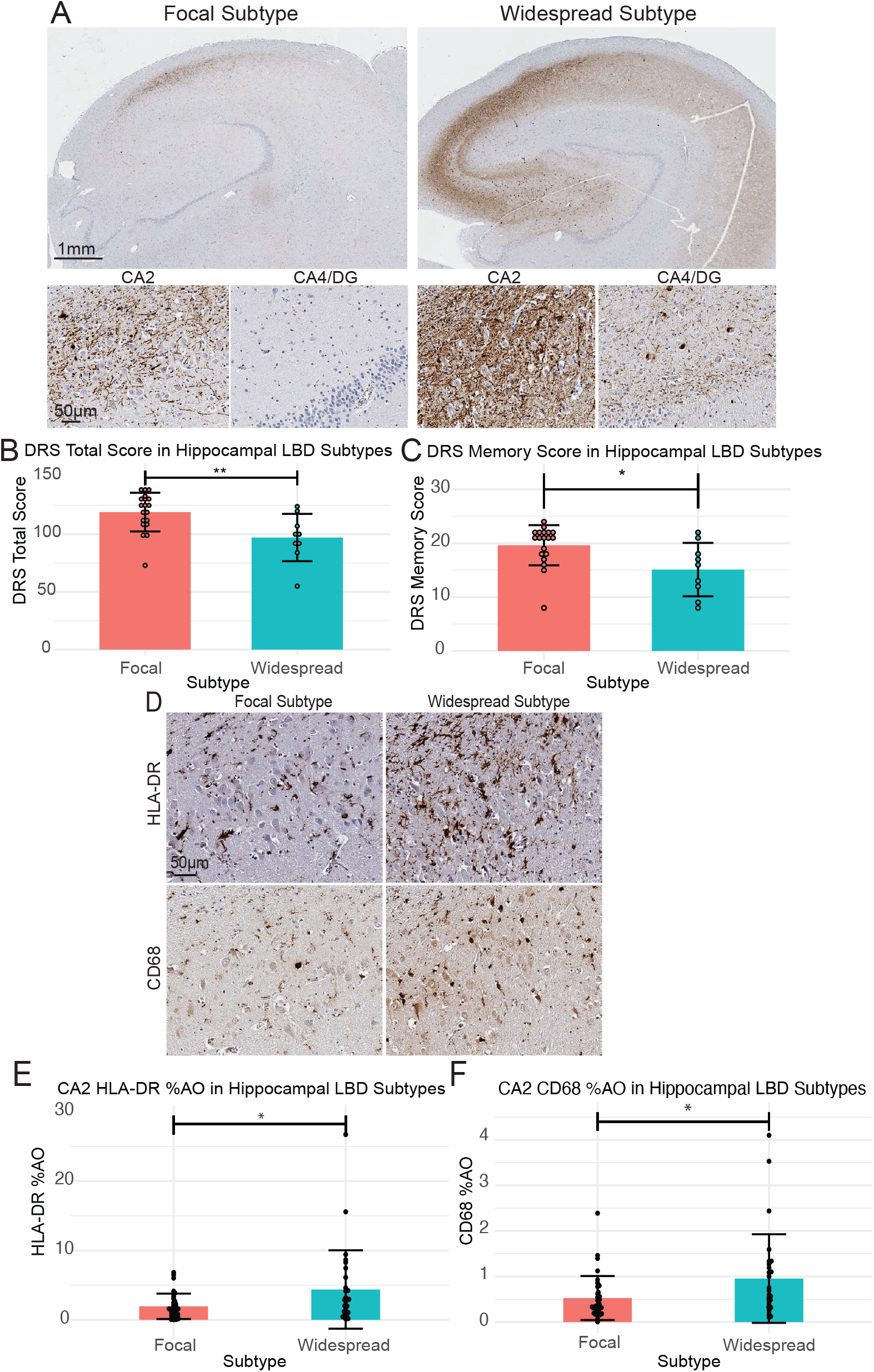
Greater n-asyn distribution in the hippocampus is associated with worse clinical outcomes and greater focal CA2 microglial reactivity. (**A**) Representative images of hippocampal sections stained for n-asyn in the identified Focal and Widespread subtypes. Insets of the CA2 and CA4/DG of each subtype are below the low power images. Scale bars indicated in images. (**B**) DRS total score and (**C**) DRS Memory score of patients whose hippocampal sections were categorized into either Focal or Widespread subtypes. Each data point represents the DRS total or DRS memory score from one patient taken at the time interval closest to death. Focal subtype *n* = 19 Widespread subtype *n* = 9. (**D**) Representative images of HLA-DR and CD68 staining in the CA2 subregion of each subtype. (**E**) Bar chart of CA2 HLA-DR %AO, (**F**) CA2 CD68 %AO in Focal vs Widespread patients. Focal subtype *n* = 34 Widespread subtype *n* = 28. All statistical analysis were done using Two tailed student T-test. All Data are mean ± SD. * p<0.05, ** p<0.01.

**Figure 5.**
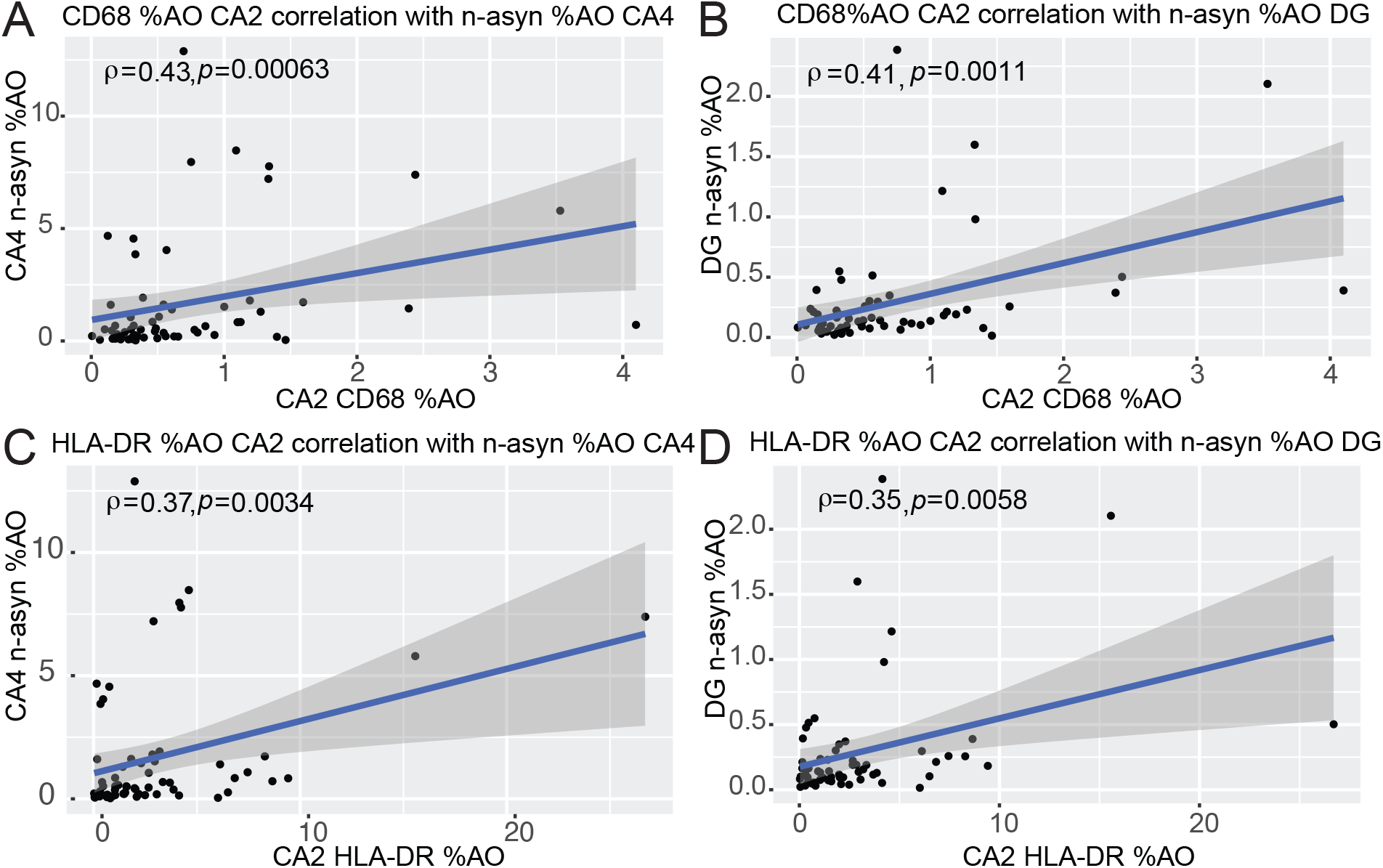
Between region correlations of focal CA2 microglial reactivity and n-asyn %AO in the CA4 and DG. (**A**) Spearman rank correlation of CA2 CD68 %AO with n-asyn %AO in CA4 and (**B**) DG. (**C**) Spearman rank correlation of CA2 HLA-DR %AO with n-asyn %AO in CA4 and (**D**) DG. Spearman rank correlation coefficient and p-values provided on charts. Blue line is best fit line, and dark grey is confidence interval.

We next asked if the measured microglial reactivity was increased in Widespread Subtype patients, and we found that Widespread Subtype patients exhibited increased HLA-DR and increased CD68 staining density compared to Focal Subtype patients, solely in the CA2 subfield (Fig. 4D-F, Supplemental Fig. 6A-J). These data illustrate that greater n-asyn levels outside of the vulnerable CA2-3 region is associated with greater microglial reactivity confined to the CA2.

**Figure 6.**
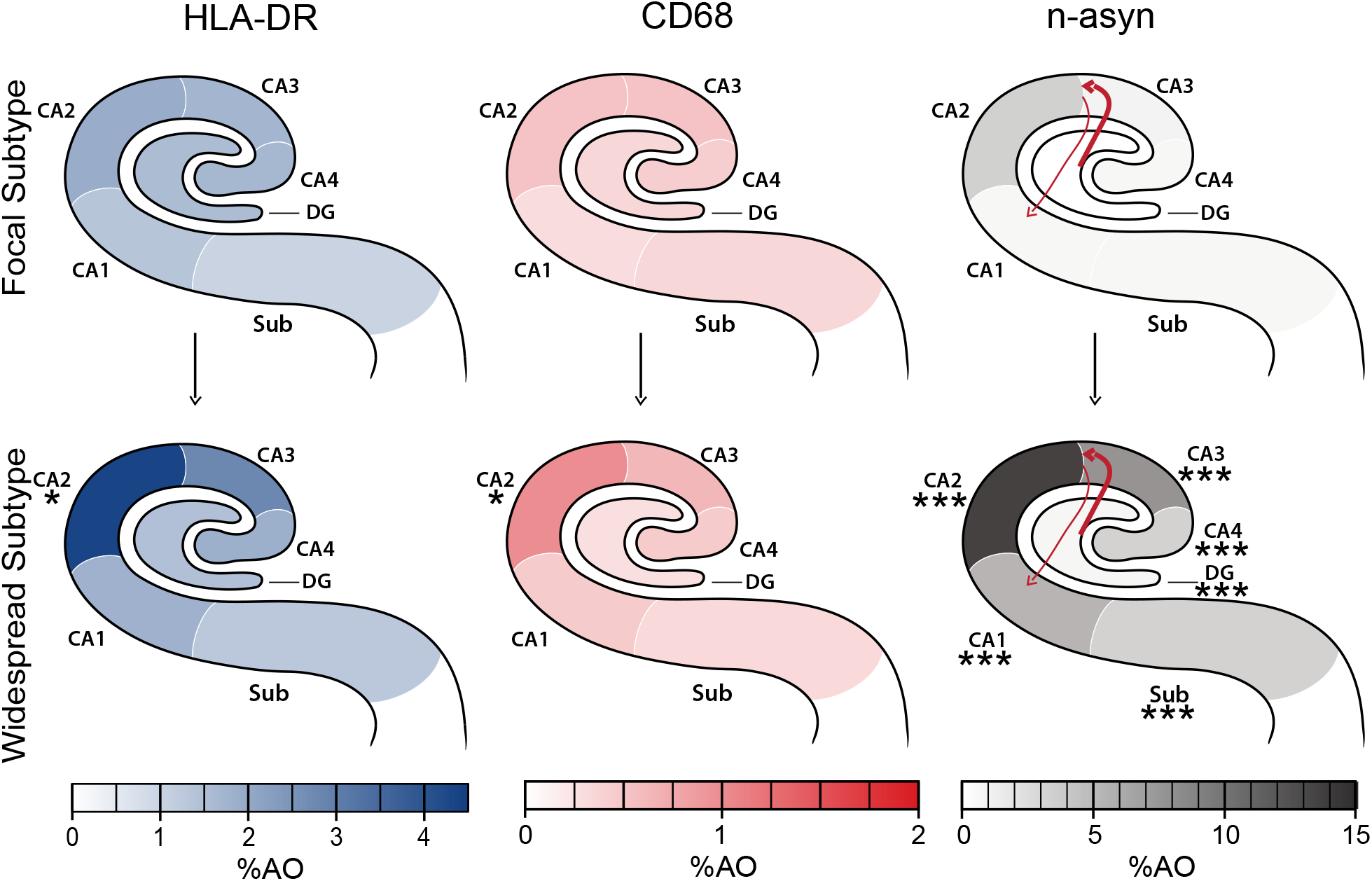
Overview schematic of modeled relative microglial reactivity and progressive n-asyn accumulation in hippocampal subregions of Focal vs Widespread subtype LBD patients. Each hippocampal schematic depicts the mean group %AO measurement for each stain (HLA-DR left, CD68 middle, n-asyn right) in each hippocampal subfield in the focal subtype (top, *n*=34) and widespread subtype (bottom, *n*=28) indicated by color intensity from the associated heat scale intensity bar below each schematic (HLA-DR=blue, CD68=red, n-asyn=grey). Asterisks denote group level difference in %AO within subregions between widespread vs focal subtypes using a Two-tailed t-test which found increased microglial reactivity to n-asyn exclusively in CA2 subfields despite greater n-asyn spread in all subfields which defined the widespread group compared to the focal group. * *p*<0.05, *** *p*<0.001. The pattern of n-asyn spread from CA2 hypothesized via anterograde and retrograde connectivity of the intrahippocampal circuit is depicted within n-asyn schematic (right), where positive correlations of CA2 CD68 and HLA-DR %AO with distal subfields with retrograde connectivity (DG, CA4) are indicated by the thicker line and lack of correlation of these markers in CA2 with subfields with anterograde connectivity (CA1, Sub) are denoted with thin line, suggesting potentially more efficient n-asyn spread via retrograde pathways associated with focal CA2 microglial activation.

### CA2 microglial reactivity is associated with n-asyn in retrograde portions of the hippocampal circuit

Despite the lack of association between microglial reactivity with n-asyn burden within each subfield outside of the CA2, we asked if the measured focal CA2 microglial reactivity correlated with n-asyn burden between hippocampal subfields that are synaptically connected via the intrahippocampal circuit. We found that both CD68 %AO and HLA-DR %AO in the CA2 correlated with n-asyn %AO in the CA4 and DG subfields (CD68 CA2 x n-asyn CA4: *ρ*=0.43, *p*<0.001; CD68 CA2 x n-asyn DG: *ρ*=0.41, *p*<0.01; HLA-DR CA2 x n-asyn CA4: *ρ*=0.27, *p*<0.01; HLA-DR CA2 x n-asyn DG: *ρ*=0.35, *p*<0.02; Fig 4A-D). n-asyn levels in neither the subiculum nor CA1 showed a correlation with the measured microglial markers in the CA2 (Supplemental Fig. 7 A-D). In the context of hippocampal connectivity, we show that focal CA2 microglial reactivity is associated with n-asyn levels in the CA4 and DG, which have a retrograde connectivity to the CA2-3 region. n-asyn %AO in the CA1 and Sub, which have anterograde connections with the CA2, did not correlate with CA2 microglial %AO.^24^ Taken together, these data suggest that focal CA2 microglial reactivity in LBD is preferentially associated with retrograde temporal progression of n-asyn through the hippocampus (Fig. 6).

## Discussion

We performed a large-scale detailed digital histopathological study using almost 300 slides from >70 autopsied patients to test the hypothesis that microglial abnormalities classically associated with a pro-inflammatory state are associated with increasing distribution and severity of n-asyn across the interconnected hippocampal subfields in LBD.^36^. Importantly, we demonstrate disease-specificity for increased microglial expression of CD68 and HLA-DR in the hippocampus in LBD, which was greatest in the n-asyn vulnerable CA2 subfield. Moreover, LBD patients with more advanced distribution of hippocampal n-asyn pathology (i.e. Widespread Subtype) have worse cognitive impairment and higher microglial reactivity relegated to the CA2 compared to LBD patients with limited hippocampal n-asyn restricted to CA2-3 region (i.e. Focal Subtype). This focal loci of microglial reactivity in the CA2 correlated with n-asyn burden in distal subfields with retrograde connectivity. These novel findings have important implications for how microglia contribute progressive n-asyn pathology in the human brain and support a potential role of modulating microglial function as a treatment strategy for LBD.

Previous studies show either no difference or a modest increase in hippocampal microglial reactivity to neuropathology compared to controls, and lower levels compared to AD.^19,20,22,23^ The reasons for these differences are likely multifactorial including various levels of mixed pathology of aging that are common in LBD and normal control cohorts and relatively limited evaluation of individual hippocampal subfields in some studies.^22,23,29^ Here, we rigorously tested for disease-specificity using a unique study design with strict neuropathological and clinical definitions to define our LBD and control cohorts to minimize other age-associated co-pathologies (Table 1). Moreover, we used multivariate modelling to account for the low-level of AD type pathology and age in our relatively “pure” LBD cohort, finding independent associations of n-asyn with the measured microglial markers (Table 2). Indeed, pathological findings in aged control cohorts are variable in terms of mixed pathology, and here, we chose to focus our control cohort to only those with PART, which is characterized by pathological tau accumulation that is relatively specific for the hippocampal CA2 subfield with minimal amyloid accumulation.^31,32,46^ LBD similarly preferentially develops n-asyn in the hippocampal CA2 subfield. Thus, our unique study design allowed us to directly test for a disease specific association of n-asyn on measured microglial abnormalities.^27^ Interestingly, we found overall similar levels of reactivity for the general microglial marker, Iba1, between the LBD and PART cohorts in all hippocampal subfields, suggesting that infiltration of microglia from surrounding brain tissue or from peripheral monocytes is not associated with n-asyn in the hippocampus or that microglia equally infiltrate the hippocampus in both cohorts. This aligns with one previous study which found similar levels of hippocampal Iba1 between PD and controls, but increased levels of CD68 expression in the CA2.^21^ In that study, there was higher relative CD68 in the CA2 compared to other brain regions in healthy controls, and there were also differences in microglial profiles between the hippocampus and substantia nigra pars compacta in PD, suggesting possible regional differences to microglial reactivity that could influence neuronal vulnerability to n-asyn.^21^ We previously found focal tau co-pathology in the CA2 associated with n-asyn in relatively pure LBD, thus we cannot completely rule out a contribution of tau pathology to hippocampal microglial reactivity.^47^ However, we did control for classical Braak tau staging in our multivariate models and included markers to two different aspects of microglial reactivity to obtain converging evidence that indicate n-asyn specifically shows a greater association with these classically associated pro-inflammatory markers relative to PART controls (Fig. 2).^34–36,48^

Next, we conducted correlational analyses to test the associations of n-asyn with the microglial markers within each subfield and found that the CA2 subfield consistently showed a moderate correlation between n-asyn and microglial reactivity but not with total Iba1 %AO (Fig. 3). These data reinforce the specific association of n-asyn in the CA2 subfield with microglial reactivity and not with a global increase in total microglia. In fact, it is increasingly recognized that microglia adopt differing states based on their local environment.^49^ Recent work in Multiple System Atrophy (MSA), a disorder characterized by the pathologic aggregation of α-synuclein within oligodendrocytes, is associated with a unique microglial subset, and our findings here raise the fascinating possibility of focal dynamic microglial states within select brain areas in LBD.^50^ Future work can examine more detailed microglial phenotyping via proteomic and transcriptomic profiles in the hippocampal CA2 subfield across LBD, AD and various aged control patients to further elucidate the microglial state specific to n-asyn, which can then be contrasted to the microglial profile associated with glial α-synuclein in MSA.

Despite the known preferential vulnerability of the hippocampal CA2 subfield to n-asyn, staging systems for LBD have limited description of hypothesized n-asyn progression within the anatomy of hippocampal subfield connectivity.^4,5,27,51^ Using quantitative digital data, we developed a limited LBD staging scheme within the hippocampal subfields to test the associations of specific microglial abnormalities with the spread of n-asyn by dichotomizing our LBD group into those with focal n-asyn pathology restricted to CA2-3 subregion (Focal Subtype) vs those with widespread n-asyn distribution (Widespread Subtype) (Figure 4). We validated this staging scheme by testing antemortem cognitive impairment between groups and confirmed that Widespread Subtype patients had worse cognitive impairment closest to death, supporting our model of advancing disease using these groupings. Interestingly, we found higher microglial reactivity associated with n-asyn selectively within the vulnerable CA2 and not within the additional subfields with n-asyn burden that defined the Widespread subtype (Figure 6). Moreover, when we specifically evaluated for associations between n-asyn and microglial reactivity between inter-connected subfields, we found that CA2 microglial reactivity had moderate correlations with n-asyn in retrograde portions of the intrahippocampal circuit (i.e. CA4 and DG) that exhibit n-asyn in our Widespread Subtype samples and correspond to worse cognitive scores (Fig. 5).^24^ In contrast, anterograde portions of the hippocampal circuit (i.e. CA1 and Sub) did not demonstrate a correlation with the measured CA2 microglial abnormalities, although these regions, also exhibit more severe n-asyn in our Widespread Subtype classification. Our human autopsy data is cross-sectional in nature, but insights from model systems can inform potential interpretation of n-asyn spread within the human hippocampus. Indeed, both *in vitro* neuronal cultures and *in vivo* murine models suggest that n-asyn-like pathology spreads more effectively through retrograde axonal transport, while anterograde transport is slow and inefficient.^52–55^ Furthermore, the injection of misfolded α-synuclein, commonly termed pre-formed fibrils, into the murine hippocampal CA4 preferentially causes n-asyn-like pathology in the CA2/3 region followed by the DG similar to what we observed here in LBD with our staging scheme.^56^ Thus, n-asyn in the CA1 and Sub could potentially spread via the less efficient anterograde mechanism or via alternative networks that are less influenced by microglial reactivity in the CA2. Integrated quantitative studies of both human autopsy and model systems can provide more comprehensive staging of n-asyn progression and microglial dynamic states to inform biological staging systems of LBD.

We performed a detailed digital histopathologic study with rigorously validated methods using multiple markers of microglial reactivity with careful multivariate modelling to account for potential confounds of aging. However, it is important to consider the potential limitations of our findings. In our unique cohort design to focus on relatively “pure” LBD patients with limited co-pathologies, we excluded participants with clinically relevant co-pathologies that may have a synergistic effect with n-asyn on microglial reactivity.^29,57^ Moreover, due to this pathologic focus on pure LBD, our cohort was enriched with clinical PD/PDD and had few dementia with Lewy body patients. Although, we accounted for co-pathologies and clinical presentation in our models, future work inclusive of mixed pathology more common in dementia with Lewy bodies can more comprehensively compare PD/PDD and dementia with Lewy body syndromes.

We focused on the hippocampus due to the well-established connectivity of the intrahippocampal circuit and well-defined neuroanatomy of the hippocampal subfields, but future studies will include larger limbic, neocortical and subcortical networks to determine if other key areas vulnerable to n-asyn share similar focal microglial reactivity and define the microglial states within those networks. It is notable that we found an association of male sex and race with an increase in both CD68 and HLA-DR signals in our LBD cohort (Table 1). Male sex has been linked to n-asyn in LBD and clinical features of LBD while ethnoracial contributions to LBD are understudied.^58–60^ Our rare autopsy cohort was from a tertiary research center and had limited ethnoracial diversity. Thus, our results may not be fully generalizable to other populations. Future work in population-based cohorts can replicate our findings and further evaluate associations of sex and ancestry with microglial reactivity in LBD.

Here we focused on a robust marker of n-asyn pathology (phosphorylated serine 129) to establish associations with microglial abnormalities, but n-asyn is well-established to adopt alternate conformations and polymorphs that can vary even within a single patient, which we did account for in this study.^40,61,62^ Furthermore, genetic variants can have alternate effects on microglial states, and future work will focus on the effects of genetic variants on microglial function in LBD, specifically.^63^ Lastly, neurodegeneration is difficult to accurately assess in the hippocampus and n-asyn may be lost with neurodegeneration.^64,65^ Although, we did not observe overt loss of n-asyn from severe neurodegeneration in our focal subtype, future experiments using reliable digital metrics of neuron loss in human tissue sections can be used to more comprehensively test associations of n-asyn, microglial reactivity, and neurodegeneration, as these are developed and validated.

Despite these limitations, we present novel data suggesting increased microglial reactivity can mark the epicenter of vulnerable brain regions in LBD and help model n-asyn progression to synaptically connected regions in the human brain. These data will have implications for biomarker studies for n-asyn and neuroinflammation in living patients, which implicate neuroinflammation in early disease.^66^ Finally, these postmortem human data provide proof-of-concept to support a therapeutic approach of modulating microglial function in LBD to slow disease progression.

## Supporting information

Combined Supplemental figures

## Data availability

All data that was generated or analysed for this publication are included as part of this publication. Data can be made available from the corresponding authors by request, except where data/results for human subjects cannot be shared due to protection of private health information.

## Acknowledgements

We would like to thank research participants who’s precious gift of brain donation has made this work possible. We thank Mary Leonard for help with the artistic design of Figure 6.

## Supplementary material

Supplementary material is available at *Brain* online

