## Supplementary material for "Microglial reactivity in the hippocampal CA2 is associated with advanced neuronal α-synucleinopathy": Combined Supplemental figures

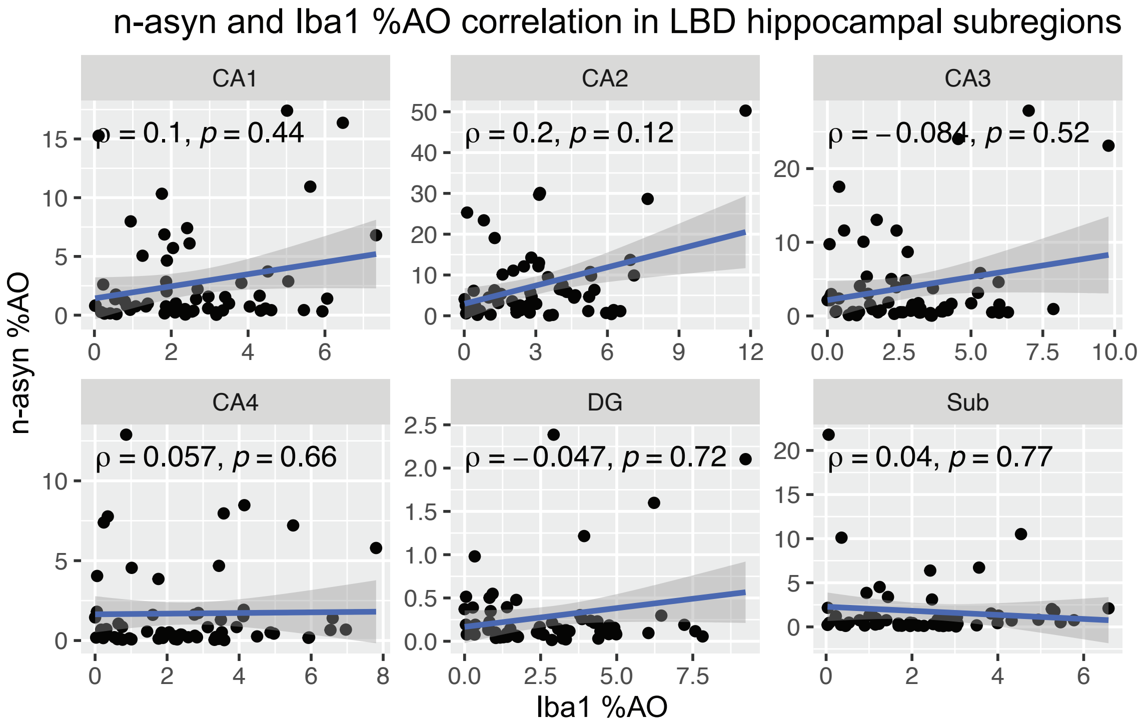


**Supplemental Figure 1 Correlation of Iba1 and n-asyn %AO in hippocampal subfields of LBD cohort.** Spearman rank correlation of n-asyn %AO in indicated subfields with Iba1 %AO. Spearman rank correlation coefficient and p-values provided on charts. Blue line is best fit line, and dark grey is the confidence interval.

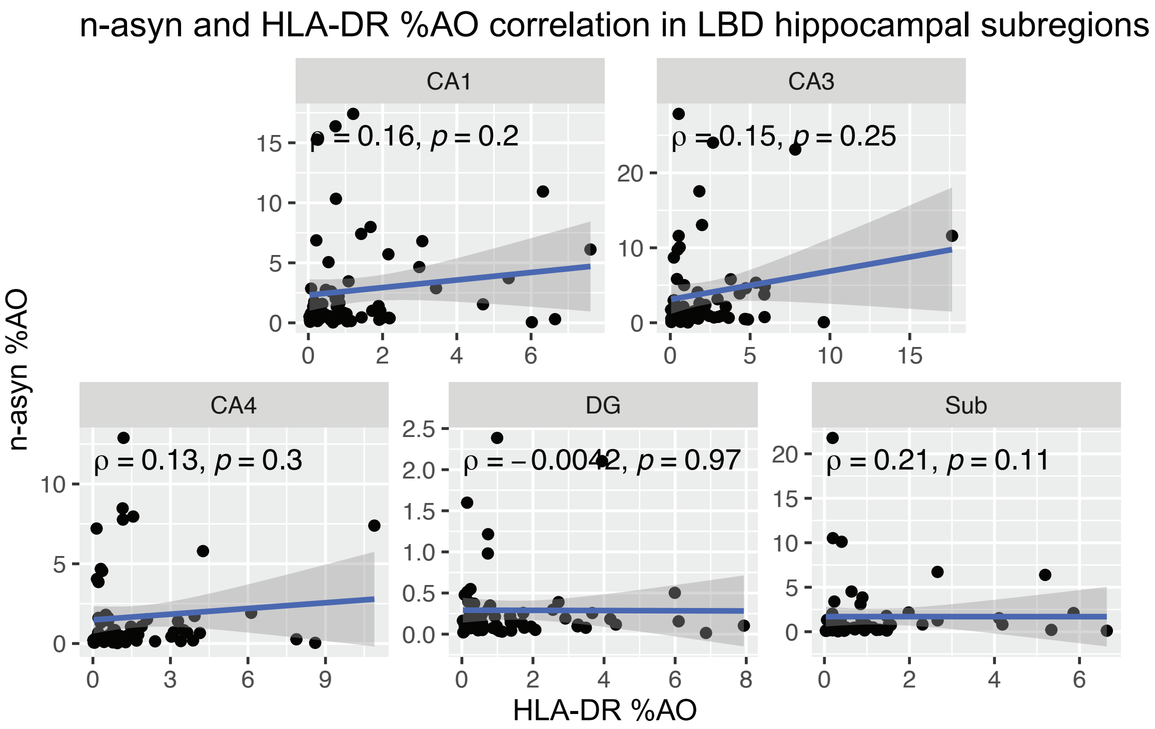


**Supplemental Figure 2 Correlation of HLA-DR and n-asyn %AO in hippocampal subfields of LBD cohort.** Spearman rank correlation of n-asyn %AO in indicated subfields with HLA-DR %AO. Spearman rank correlation coefficient and p-values provided on charts. Blue line is best fit line, and dark grey is the confidence interval.

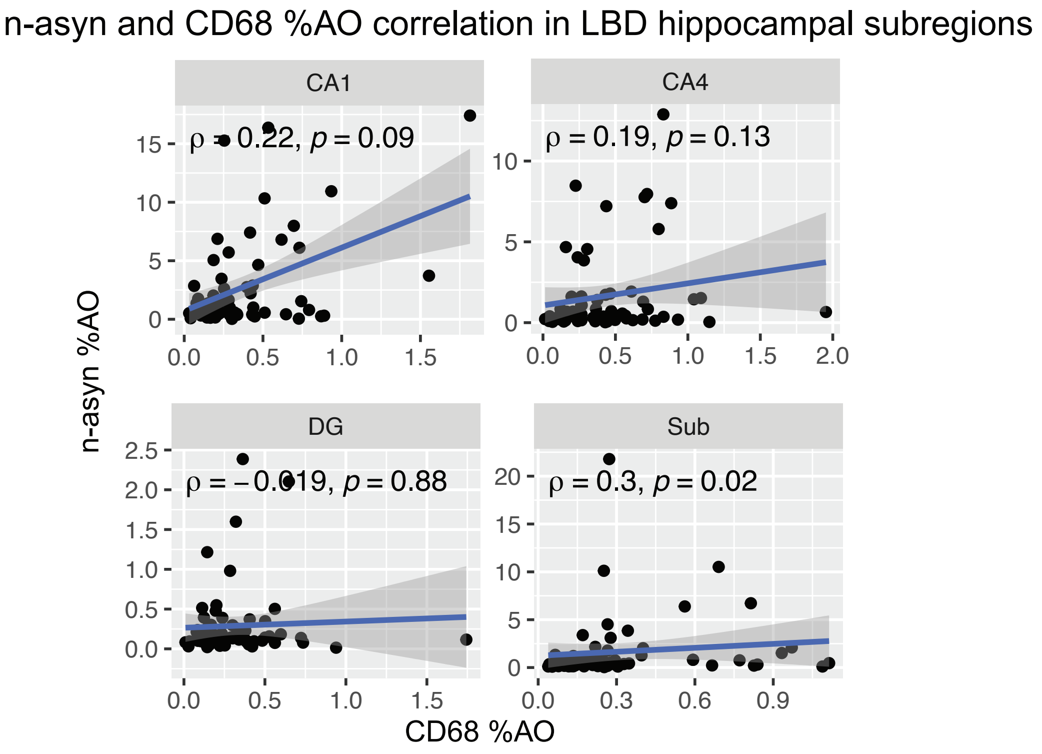


**Supplemental Figure 3 Correlation and CD68 and n-asyn %AO in hippocampal subfields of LBD cohort.** Spearman rank correlation of n-asyn %AO in indicated subfields with CD68 %AO. Spearman rank correlation coefficient and p-values provided on charts. Blue line is best fit line, and dark grey is the confidence interval.

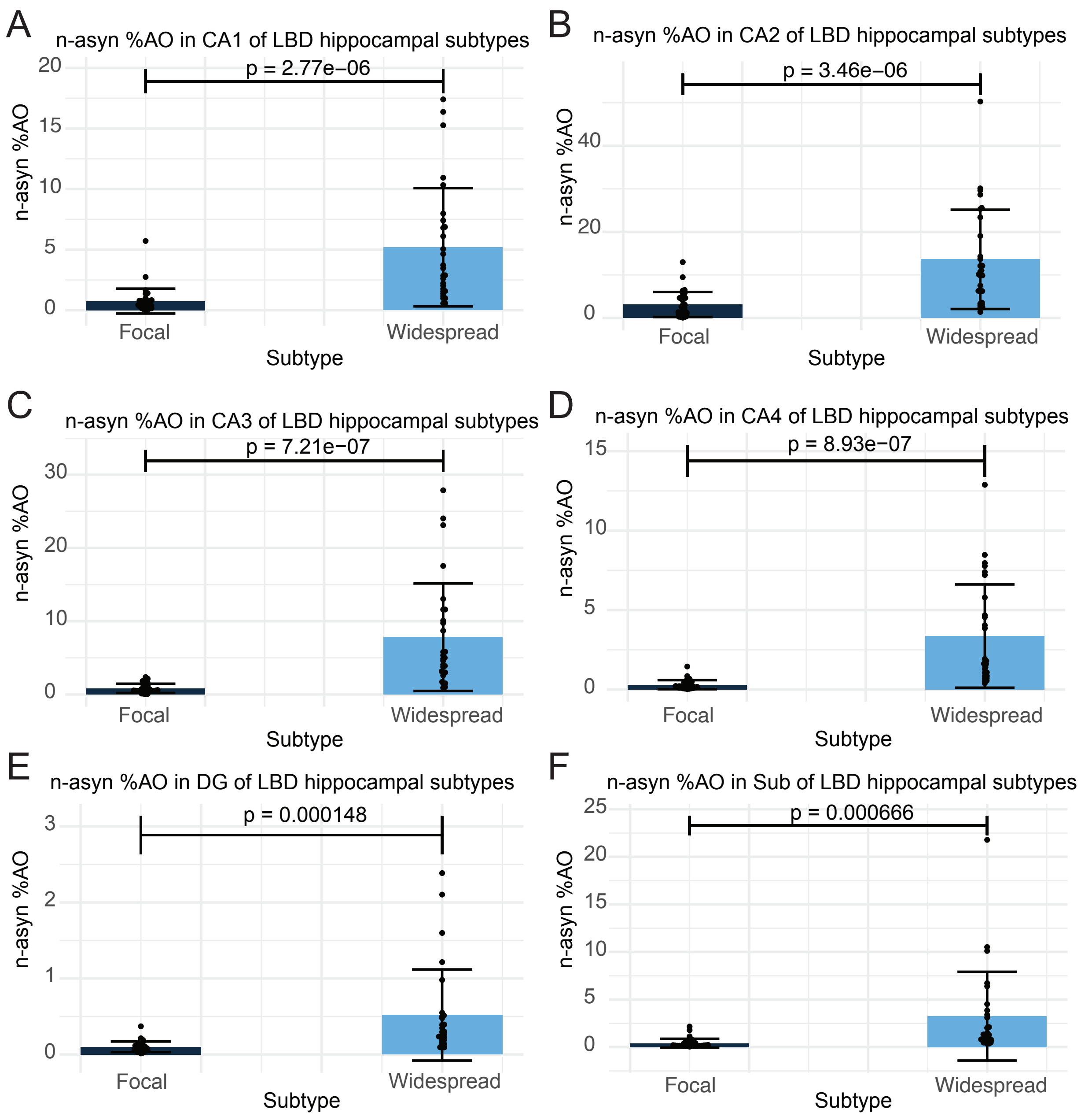


**Supplemental Figure 4 Widespread subtype patients exhibit increased n-asyn %AO in all hippocampal subfields.** (A-F) Bar charts of n-asyn %AO in Focal subtype n = 34 and Widespread subtype n = 28 in indicated hippocampal subfields. Two tailed student T-test of n-asyn %AO between patient groups. All Data are mean ± SD. p-values indicated on charts.

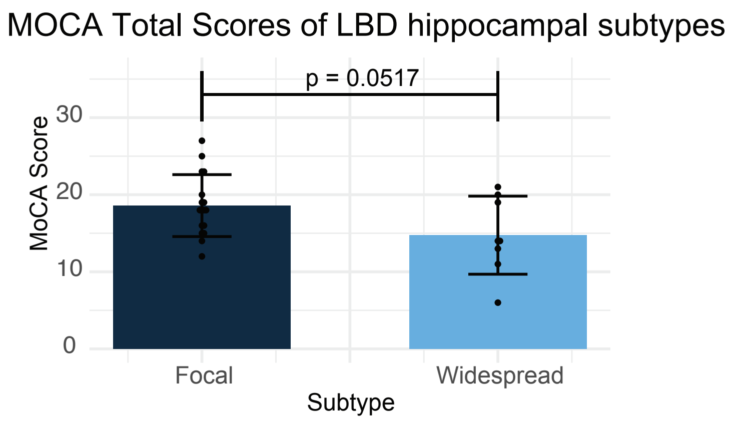


**Supplemental Figure 5 Comparison of MOCA scores between Focal and Widespread subtype patients**. Bar chart of MOCA scores in Focal subtype n = 17 Widespread subtype n = 8. Two tailed student T-test of MOCA Total Scores between patient groups. All Data are mean ± SD. p-value indicated on charts.

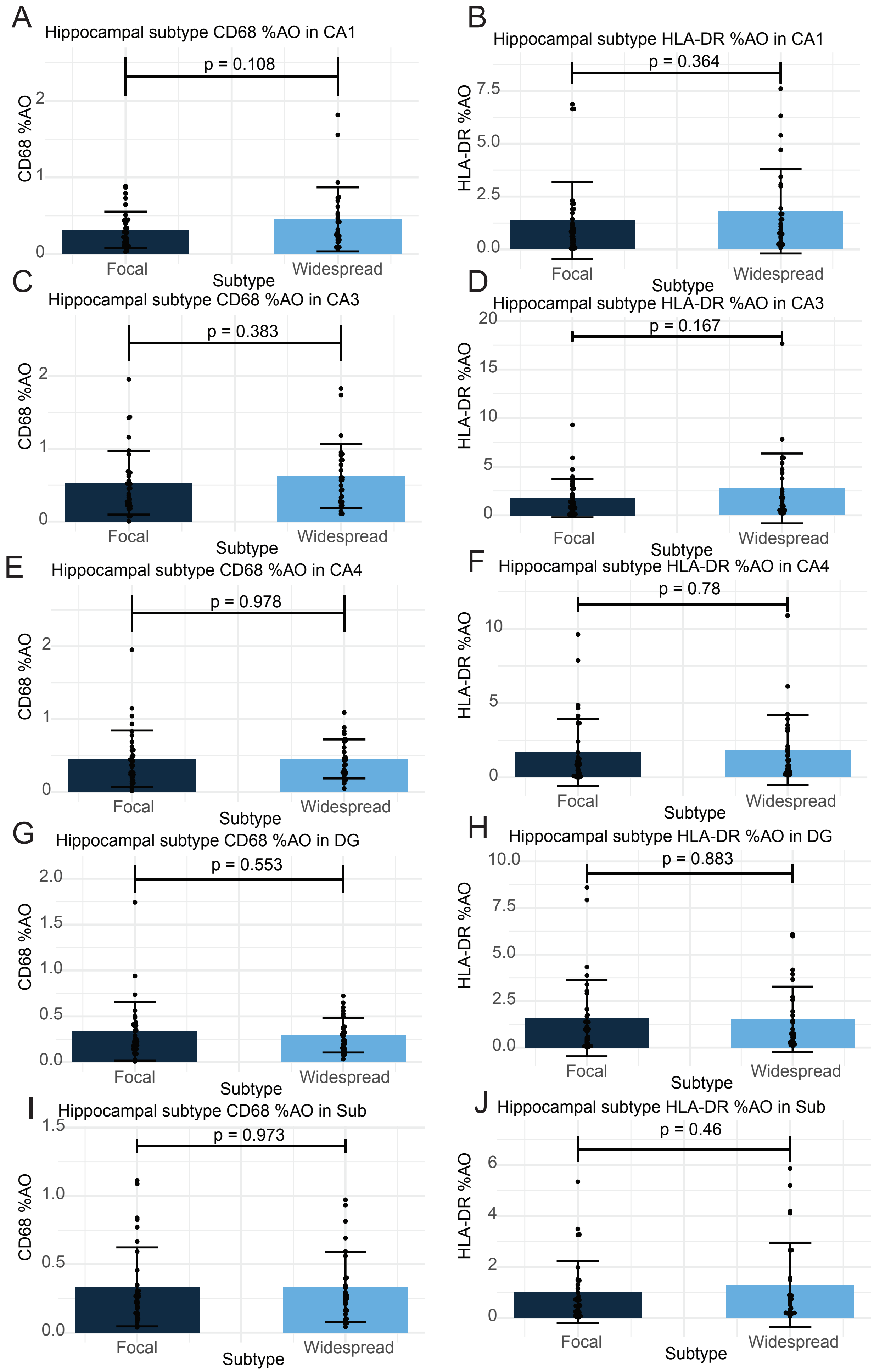


**Supplemental Figure 6 There is no difference in microglial reactivity in any hippocampal subregion outside the CA2 between Focal and Widespread subtype patient groups.** (A-J) HLA-DR and CD68 %AO within each hippocampal subregion outside the CA2, indicated on charts, in Focal and Widespread subtype patient groups. in Focal subtype n = 34 and Widespread subtype n = 28. Two tailed student T-test. All Data are mean ± SD. p-value indicated on charts.

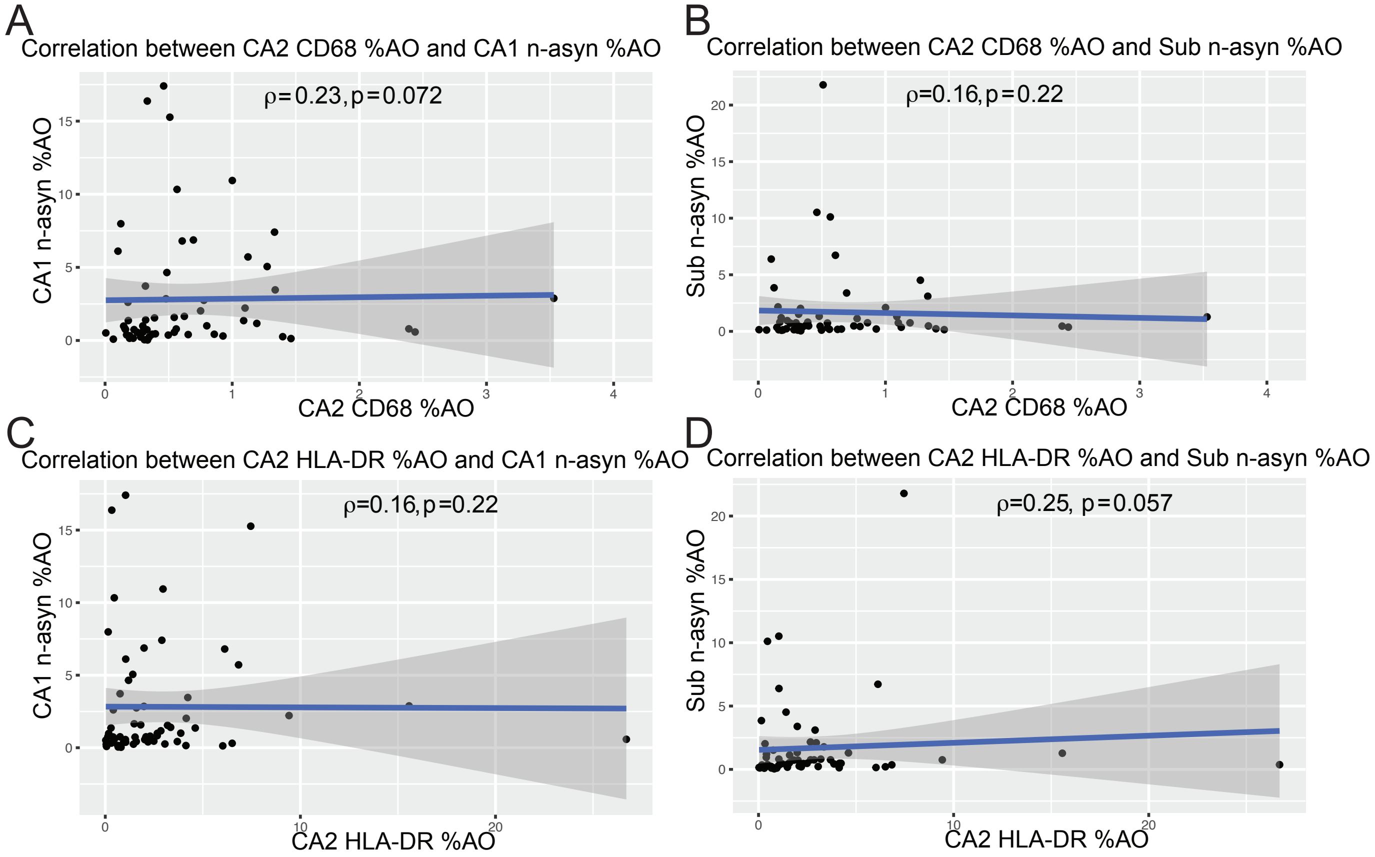


**Supplemental Figure 7 Between region correlations of focal CA2 microglial reactivity and n-asyn %AO in anterograde portions of the intrahippocampal circuit.** (A) Spearman rank correlations of CA2 CD68 %AO with n-asyn %AO in CA1 and (B) Sub. (C), Spearman rank correlations of CA2 HLA-DR %AO with n-asyn %AO in CA1 and (D) Sub. Spearman rank correlation coefficient and p-values provided on charts. Blue line is best fit line, and dark grey is confidence interval.
